# Identification of MYC synthetic lethal genes and networks

**DOI:** 10.1101/2024.04.25.590465

**Authors:** Timothy D. Martin, Mei Yuk Choi, Rupesh Patel, Anthony Liang, Mamie Z. Li, Stephen J. Elledge

## Abstract

MYC is a potent oncogene that is frequently overexpressed in human tumors arising in different tissues. To date there are no approved therapies to directly antagonize oncogenic MYC and its role in driving tumorigenesis. As an alternative approach we employed genetic screens using CRISPR and shRNA to identify the genes that are required for the survival and growth of cells harboring high levels of MYC expression. We find that cells with elevated MYC require the expression of many pro-growth and metabolic pathways including genes involved in mitochondrial citrate production and transport. This citrate producing pathway is critical for cells with elevated MYC to generate the necessary acetyl-CoA to drive the lipid synthesis required for increased proliferation. Inhibition of this pathway results in reduced proliferation and in vivo tumor growth providing a potential therapeutic strategy to target MYC-driven cancers.

**HIGHLIGHTS:**

– CRISPR and shRNA screens identify synthetic lethal interactions with overexpressed MYC
– MYC overexpressing cells are more sensitive to disruption of citrate production and transport
– Inhibition of SLC25A1 reduces growth of MYC driven tumors

## INTRODUCTION

The myelocytomatosis or MYC protein was originally described as an oncogene (*v-MYC*) acquired by transformed cells infected with the avian leukemia virus MC29 (Watson et al., 1983). Subsequent studies found that the cellular counterpart, c-MYC (hereafter termed MYC), is overexpressed in many human tumor types where gene amplification of *MYC* occurs in ∼21% of human tumors and predicts poorer overall survival (Schaub et al., 2018). The MYC protein itself can be stabilized through mutation or deletion of the tumor suppressor and MYC E3 ligase FBXW7 and oncogenic activation of the Ras pathway results in enhanced MYC stability through ERK phosphorylation of MYC on serine 62 (Sears et al., 2000; Welcker et al., 2004; Yada et al., 2004). MYC functions as an E-box transcription factor that forms a heterodimer with its partner Max to regulate gene expression and drive cells into S phase (Topham et al., 2015). MYC can also heterodimerize with its partner Miz to repress transcription and prevent senescence (Wanzel et al., 2003). In tumors, increased MYC expression is thought to allow for MYC binding at lower affinity sites in promoters and enhancers leading to an amplification of transcriptional output that drives cell growth (Lin et al., 2012). Intense research into the role of MYC in cancer biology has shown that high levels of MYC expression is sufficient for spontaneous tumorigenesis and MYC-driven tumors require sustained high MYC expression for growth (Felsher and Bishop, 1999; Stewart et al., 1984). This increase in MYC works both directly and indirectly to enhance the expression of target genes many of which promote growth through upregulation of anabolic pathways including nucleotide, lipid, and protein biosynthesis (Dang et al., 2009; Stine et al., 2015). The ability to target MYC and reduce its pro-tumor activity has been elusive though recent promising pre- clinical inhibitors have been described (Han et al., 2019). Other MYC targeting approaches include expression of a MYC-binding helix termed Omomyc that can sequester MYC to prevent transcriptional activation and KRAS mutant lung tumors are sensitive to Omomyc expression (Beaulieu et al., 2019; Soucek et al., 1998).

An alternative approach to directly inhibiting MYC is to identify the genes that are required specifically in the context of MYC overexpression. Previous work by our group and others has identified a number of these genes using RNAi based screens (Kessler et al., 2012; Liu et al., 2012; Toyoshima et al., 2012). These identified MYC synthetic lethal (SL) targets are genes that are dispensable for the survival of cells with endogenous levels of MYC but are required for the growth and survival of cells that have high levels of MYC like those seen in cancer (Cermelli et al., 2014). This feature makes SL genes attractive therapeutic targets since they are less required for normal cell survival and growth and are more likely to generate tumor specific responses. The concept of synthetic lethality and cancer therapy is perhaps most notable in the profound sensitivity of BRCA-deficient cells to PARP inhibition (Bryant et al., 2005; Farmer et al., 2005).

Previous synthetic lethal studies have demonstrated a reliance of oncogene expressing cells on cellular stress response pathways in what is known as non-oncogene addiction (NOA) (Luo et al., 2009b). NOA occurs due to the increased demands of cells expressing an oncogenic driver on key cellular processes. These could be general aspects of cellular homeostasis such protein translation, the DNA damage response, or mitosis that are stressed to keep up with increased cellular proliferation or due to an imbalance of some sort caused by inappropriately hyperactivating an oncogenic pathway in the absence of the normal developmental cues that normally accompany it. For example, oncogenic KRAS mutant cells are more reliant on mitotic fidelity and perturbing mitosis either genetically or pharmacologically leads to increased toxicity that can be exploited to decrease tumor growth (Luo et al., 2009a). This is in part due to KRAS activation leading to mitotic infidelity. Previous studies also show that cells with elevated MYC expression have a dependency on a SUMOylation-dependent transcriptional network that is required for proper cell division and a reliance on tRNA synthetases for increased proliferation (Kessler et al., 2012; Zirin et al., 2019). While these aren’t traditional synthetic lethal interactions, therapeutically exploiting various non-oncogene addiction pathways including proteostasis and nucleotide metabolism results in reduced tumor growth (Dai et al., 2007; Gad et al., 2014; Huber et al., 2014).

In this study, we used both genome-wide CRISPR and shRNA essential gene libraries to identify candidate MYC SL genes. We found that cells with elevated MYC expression are more sensitive to loss of metabolic pathways including genes involved in nucleotide, lipid, and protein biosynthesis. Inhibition of stress response networks like the DNA damage and unfolded protein responses (UPR) cause increased toxicity in cells with high MYC expression representing a therapeutically targetable type of non-oncogene addiction whose inhibition can impede MYC amplified cell growth. Interestingly, we show that cells with elevated MYC are dependent on the mitochondrial production of citrate via citrate synthase and its subsequent transport to the cytosol through the transport protein SLC25A1. Cytosolic citrate is then available for acetyl-CoA production through the action of ACLY which generates acetyl-CoA for lipogenesis, a necessity for the increased cell growth demands caused by MYC overexpression. Inhibiting SLC25A1 results in reduced proliferation and *in vivo* MYC-driven tumor growth providing a new therapeutic target to combat tumors harboring high levels of MYC expression.

## RESULTS

### CRISPR and shRNA-based proliferation screens to identify synthetic lethal interactions with overexpressed MYC

We generated human colonic epithelial cells (HCECs) with a doxcycycline (dox) inducible MYC transgene driven by a tetracycline response element (TRE) promoter (TRE-MYC). Treatment of these cells with dox results in high levels of MYC protein expression (Figure 1A). In these cells, we introduced a lentiviral genome-wide CRISPR library to identify synthetic lethal (SL) genes (Figure 1B). After introduction of the library, cells were treated with DMSO or dox to upregulate MYC and allowed to grow for ten additional population doublings (PDs). Cells treated with dox to increase MYC expression exhibited increased proliferation and sgRNA abundance was measured at both PD5 and PD10 to compare to an input PD0 timepoint (Figure 1C). Analysis of the screening results indicated a dox-independent reduction in sgRNAs targeting essential genes like ribosomal proteins and chaperones like HSPE1 (Figure 1D and Supplemental Tables 1 and 2). We also observed an increase in sgRNA abundance for known growth suppressing genes like TP53 (Figure 1D). To identify SL genes we analyzed the data to find genes whose sgRNAs significantly depleted in the MYC High (+ dox) condition but had a limited effect on the MYC WT (no dox) cells, some of which are labeled in Figure 1D. We curated a list of 314 putative MYC SL genes by identifying genes that were at least 2-fold more toxic with an FDR less than 0.3 in MYC High cells compared to MYC WT (Supplemental Table 2). We performed gene set enrichment analysis (GSEA) using Hallmark and KEGG gene sets to identify pathways and terms shared by these MYC synthetic lethal genes (Figure 1E). Of note these included known MYC target genes, cell cycle related genes, metabolic pathways including the TCA cycle, and stress response pathways like the unfolded protein response (UPR) (Figures 1E and S1A). Western blotting of the TRE-MYC HCECs indicated an increase in ER Stress and UPR related proteins ATF-4 and CHOP and MYC High expressing cells were more sensitive to the ER stress and UPR inducer tunicamycin (Figure S1B and S1C).

**Figure 1.**
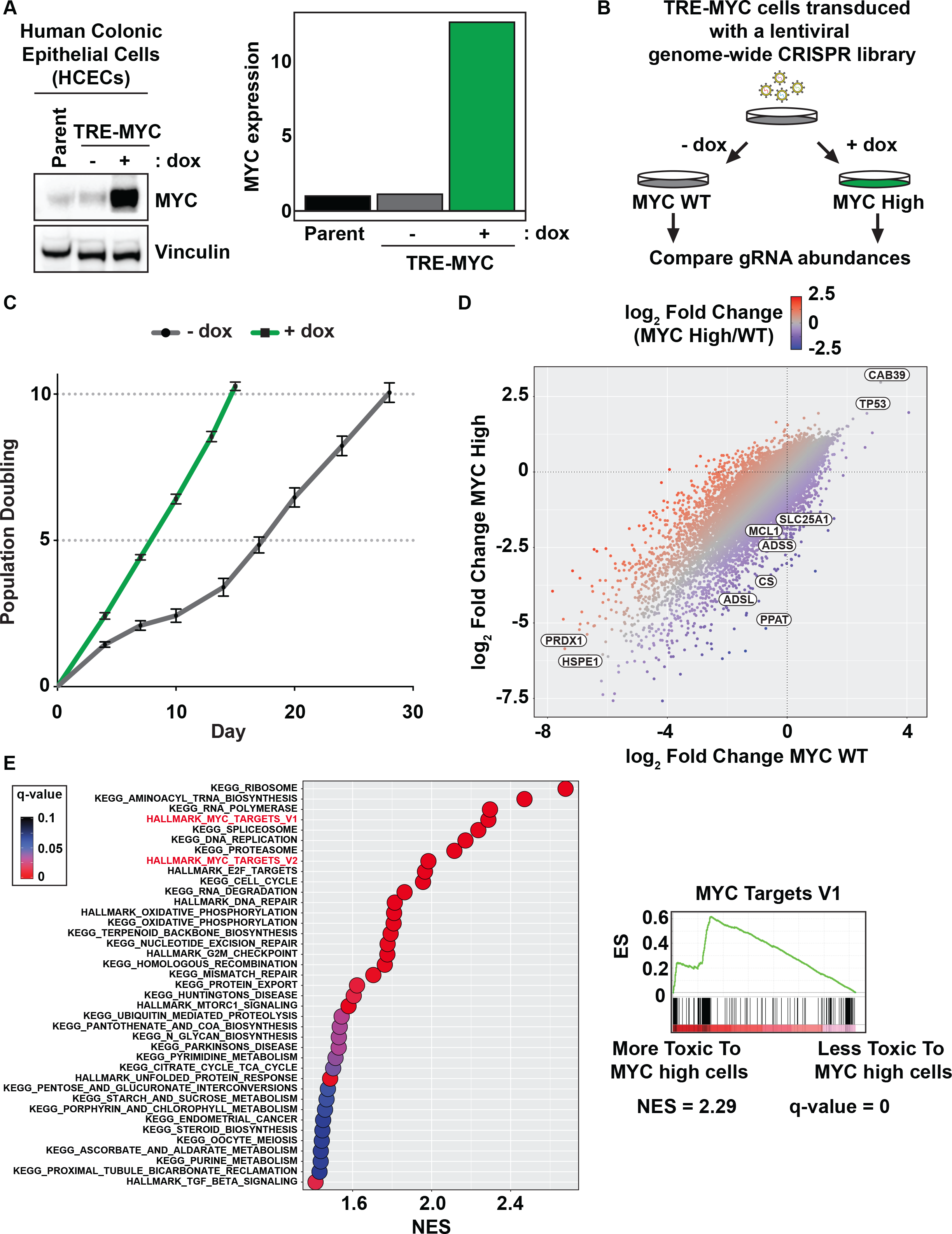
A genome-wide CRISPR screen to identify synthetic lethal (SL) interactions in human colon cells overexpressing the oncogene MYC. A. Doxycycline regulated overexpression of MYC in human colonic cells (HCECs). HCECs were infected with a pInducer20-based lentivirus containing TRE-MYC UbC-rtTA3-IRES- blast to generate the TRE-MYC cell line. Blasticidin resistant cells were examined for MYC expression after 48 h of 100 ng/mL doxycycline treatment. Bar graph represents the fold induction of MYC before and after dox treatment as determined by LI-COR quantification of the western blot. B. Schematic for the MYC SL CRISPR screen. Cells were infected in triplicate with a genome-wide lentiviral CRISPR library, selected with puromycin, and then split into minus and plus dox conditions. Sequencing of the sgRNAs was performed on the initial infected population and after 5 and 10 population doublings (PDs) with and without 100 ng/mL dox treatment. **C.** Inducing MYC overexpression with dox results in increased proliferation in HCECs. Growth curves from the CRISPR screen described in panel B. **D.** Comparison of the CRISPR screen results. The log2 fold change for each gene in the CRISPR library was plotted for the plus dox and minus dox conditions after PD10. Loss of essential genes (PRDX1 & HSPE1) is universally toxic while loss of potent tumor suppressors (TP53 & CAB39) enhances cell growth regardless of MYC status. Genes that are more toxic to cells with high levels of MYC (PPAT, SLC25A1, MCL1, etc) are candidate MYC SLs. **E.** Gene set enrichment analysis identifies MYC SL pathways. The log2 fold change differential between the minus and plus dox arms of the CRISPR screen were used to rank genes. This gene ranking was used to identify MYC SL pathways using GSEA.

Next we analyzed transcriptome changes by performing RNA-seq in the TRE-MYC cells during MYC WT and MYC High conditions (Figure S1D). MYC overexpression increased the expression of known MYC target genes and growth promoting pathways as seen in GSEA (Figure S1E and Supplemental Table 3). We compared our list of candidate MYC SL genes to the transcriptome data and found no statistical correlation indicating that one cannot predict a gene’s potential genetic dependency when MYC is overexpressed by an elevation in its transcription upon MYC overexpression (Figure S1F).

Essential genes are universally toxic in CRISPR screens but hypomorphic effects are still applicable to developing therapies. The fact that inhibition of a gene is toxic does not mean that it cannot be used therapeutically. Many drugs targeting otherwise essential proteins are under clinical evaluation including drugs targeting the essential mitotic kinase PLK1 and MYC-driven multiple myelomas are sensitive to RNA polymerase I inhibition which limits ribosomal biogenesis (Gutteridge et al., 2016; Lee et al., 2017). To explore the role of essential genes in MYC biology, we generated an improved shRNA library using the SPLASH algorithm that targets ∼1,100 essential genes to further uncover MYC SL genes (Pelossof et al., 2017). Since the shRNA essential gene library is doxycycline inducible, we made HCECs stably expressing high levels of MYC constitutively from an EF1α promoter (Figure S2A). We allowed MYC WT and MYC High cells to proliferate for 10 PDs after shRNA induction and analyzed shRNA abundance at PD3, PD7, and PD10 as compared to PD0 input (Supplemental Table 4). Many essential genes including ribosome and proteasome subunits showed increased toxicity when depleted in MYC High cells (Figure S2B). Depletion of 6-phosphogluconate dehydrogenase (PGD), an early enzyme in the pentose-phosphate pathway, was toxic to MYC High cells. We individually cloned a panel of the shRNAs from the SPLASH library targeting PGD and used these shRNAs to deplete PGD expression in MYC WT and MYC High cells (Figure S2C). Proliferation assays showed that genetic inhibition of PGD can reverse the increase in proliferation seen with MYC overexpression to levels comparable to MYC WT cells (Figure S2D).

### Genetic and pharmacological validation of candidate MYC synthetic lethal genes

From our CRISPR screening results we generated a list of the top MYC SL genes which are displayed in Figure 2A. These are the genes whose depletion caused the biggest decrease in MYC High cell growth without significantly limiting the growth of MYC WT cells. To validate these candidates as MYC SL genes we used a multicolor competition assay (MCA) where non-fluorescent MYC High and GFP-positive MYC WT cells are mixed and changes in the cell populations are monitored by flow cytometry after depletion or inhibition of candidate MYC SL genes (Figure 2B). Depletion of a MYC SL gene will reduce the ability of MYC High cells to outcompete the MYC WT cells during the assay resulting in an observed increase in fluorescent cells in the population compared to controls. One candidate MYC SL gene is the anti-apoptotic protein MCL1, an inhibitor of BAX/BAK whose haploinsufficiency has been previously shown to protect mice from MYC-driven leukemia (Xiang et al., 2010). Expression of Cas9 and sgRNAs targeting MCL1 increases the abundance of GFP+ MYC WT cells compared to MYC High indicating a toxicity to MYC High cells (Figure 2C). Treatment with the MCL1 inhibitor S63845 had a similar effect and MYC High cells had a reduced GR50, the concentration that gives half maximal proliferation inhibition using normalized growth rates, compared to MYC WT cells (Figures 2D and S3A) (Hafner et al., 2016). MYC High cells were visually apoptotic after treatment with S63845 while MYC WT cells showed no apoptosis induction as measured by cleaved caspase-3 and cleaved PARP (Figures S3B and S3C).

**Figure 2.**
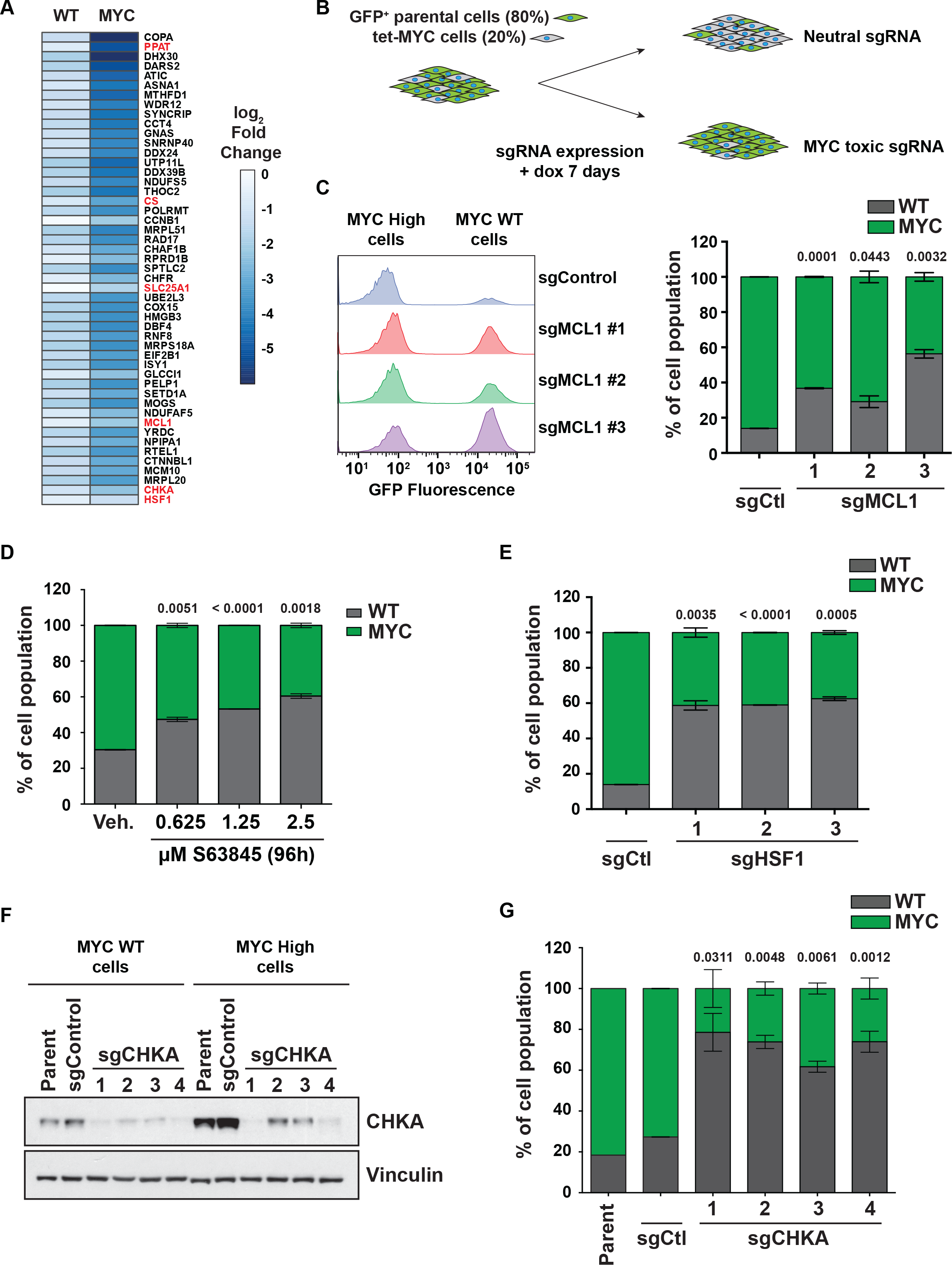
Validation of a panel of MYC synthetic lethal genes. A. A heatmap showing some of the top scoring MYC SL genes from the CRISPR screen. Gene names in red are validated in this figure or accompanying figures. **B.** Diagram of the multicolor competition assay (MCA) to validate MYC SL genes. Non-fluorescent TRE- MYC cells were mixed with GFP expressing parental cells at a 20%:80% (TRE- MYC:parental) ratio. Cell mixtures were then infected with Cas9/MYC SL gene sgRNA vectors, selected with puromycin, treated with 100 ng/mL doxycycline, and allowed to proliferate for 1 week. After 1 week, cell mixtures were examined by flow cytometry to determine the ratio of TRE-MYC:parental cells. **C.** MCL1 is a MYC SL gene. Representative flow cytometry and MCA data from 3 MCL1 sgRNAs and an AAVS1 targeting control. Disruption of MCL1 reduces the growth of MYC overexpressing non-fluorescent cells resulting in an increase in GFP+ WT MYC contribution to the overall cell population. **D.** Small molecule inhibition of MCL1 is more toxic to MYC overexpressing cells. Cells were mixed as in panel B and treated with the MCL1 inhibitor S63845 for 4 days. Cell mixtures were examined by flow cytometry after 4 days. **E.** The transcription factor HSF1 is a MYC SL gene. Cells were mixed as in panel B and infected with HSF1 or an AAVS1 control sgRNA. Following 1 week, cell mixtures were examined by flow cytometry. **F.** Choline kinase (CHKA) targeting using CRISPR. TRE-MYC cells were infected with CHKA targeting or AAVS1 control sgRNAs. Cells were treated with vehicle or 100 ng/mL dox for 7 days and cell lysates were western blotted with the indicated antibodies to determine CHKA protein disruption. **G.** Choline kinase (CHKA) depletion is toxic to MYC High cells. Cells were mixed as in panel B and infected with CHKA or an AAVS1 control sgRNA. Following 1 week, cell mixtures were examined by flow cytometry. Numbers above bar graphs indicate the p- value from an unpaired Student’s t-test to compare sgRNA or treatment groups to sgControl or Vehicle, respectively.

Other candidate MYC SL genes included choline kinase alpha (CHKA), HSF1, and PPAT (Figure 2A). CRISPR targeting of HSF1, a heat shock inducible transcription factor, resulted in reduced growth of MYC High cells as measured by MCA (Figures 2E and S3D). CRISPR depletion of CHKA, a kinase involved in phospholipid synthesis, showed similar effects (Figure 2F and 2G). Inhibition of CHKA using the inhibitor MN58b also reduced MYC High cell proliferation (Figure S3E). PPAT, a gene involved in nucleotide biosynthesis and a direct MYC target gene, was targeted using CRISPR and also found to be required for MYC High cell proliferation (Figure S3F).

### Mitochondrial citrate production and transport are required to support overexpressed MYC induced proliferation

Analysis of our CRISPR and shRNA screening data identified genes (SLC25A1, CS, and ACLY) involved in citrate and acetyl-CoA metabolism as candidate MYC SL genes (Figure 3A). MYC High cells are known to rely upon glutamine to replenish TCA cycle intermediates including citrate to drive metabolic growth (Gao et al., 2009). Citrate generated in the TCA cycle, by citrate synthase (CS) or through the reductive carboxylation of α-ketoglutarate by IDH enzymes (Metallo et al., 2011), can be exported from the mitochondria by SLC25A1/CTP to the cytosol where cleavage by ACLY generates acetyl-CoA that is then available for fatty acid and sterol synthesis as well as histone and protein acetylation (Figure 3B). We stably expressed Cas9 and sgRNAs targeting SLC25A1 or CS in TRE-MYC HCECs. Cell lysates were western blotted to confirm disruption of SLC25A1 and CS protein expression (Figure 3C and 3D). To determine changes in proliferation, these same sgRNAs were used in MCAs as described earlier in Figure 2B. For both SLC25A1 and CS, CRISPR targeting led to a reduction in MYC High cell proliferation (Figure 3E and 3F). Analysis of the Cancer Dependency Map (DepMap) revealed that colorectal tumor cell lines with MYC gene amplification are more sensitive to SLC25A1 targeting by CRISPR compared to cell lines that have two copies of MYC (Figure 3G).

**Figure 3.**
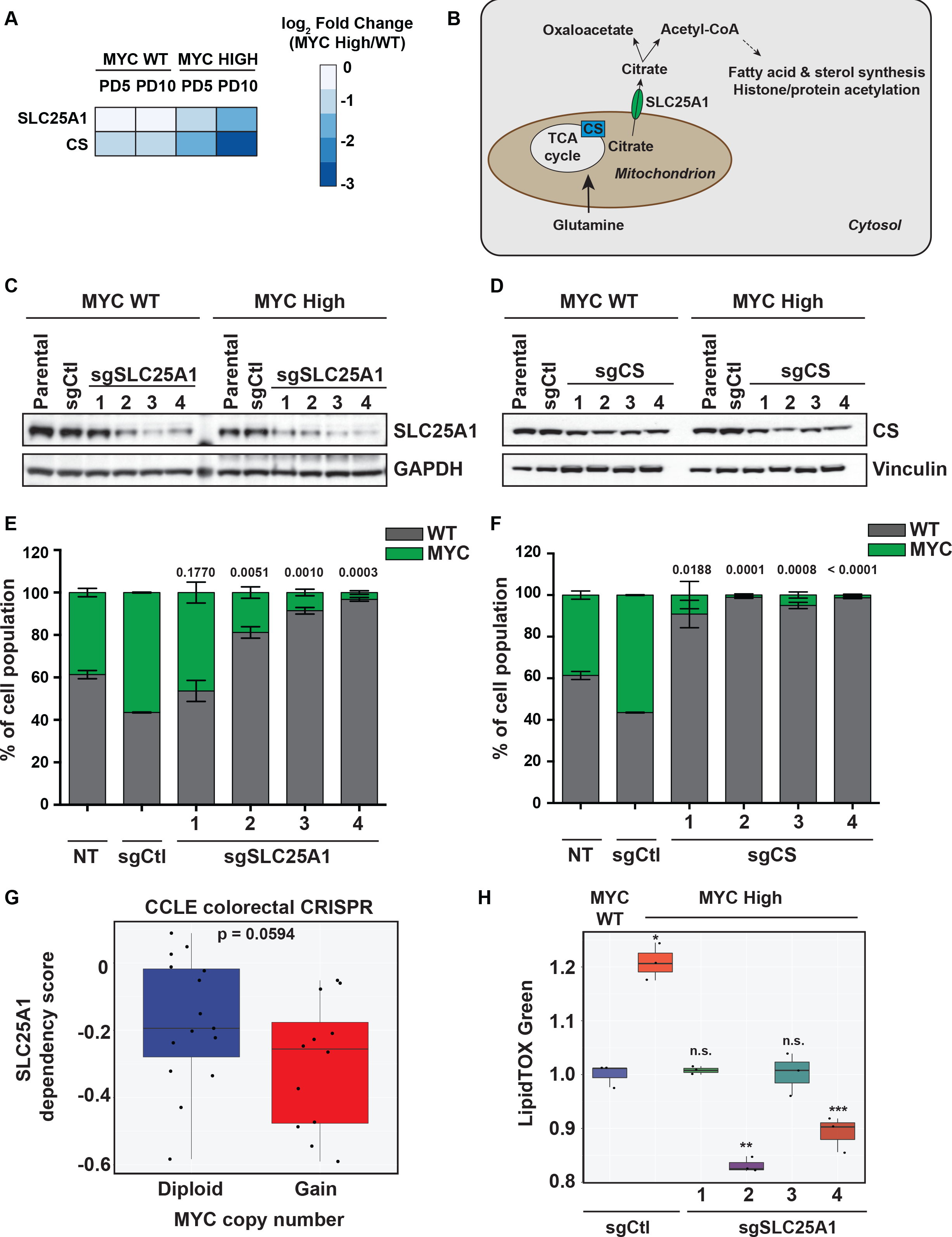
Disruption of mitochondrial citrate production and transport is toxic to MYC overexpressing cells. A. The citrate transporter SLC25A1 and citrate synthase (CS) are MYC SL genes. A heatmap of CRISPR screening results showing the log2 fold changes of SLC25A1 and CS at PD5 and PD10 in MYC WT and MYC overexpressing cells. **B.** Schematic of SLC25A1 and CS function. Mitochondrial citrate generated in the TCA cycle by CS can be shuttled to the cytosol by SLC25A1 where it can be utilized to generate acetyl-CoA, an important step in fatty acid synthesis and protein/histone acetylation. **C.** SLC25A1 targeting using CRISPR. TRE-MYC cells were infected with SLC25A1 targeting or AAVS1 control sgRNAs. Cells were treated with vehicle or 100 ng/mL dox for 7 days and cell lysates were western blotted with the indicated antibodies to determine SLC25A1 protein disruption. **D.** CS targeting using CRISPR. TRE-MYC cells were infected with CS targeting or AAVS1 control sgRNAs. Cells were treated with dox as in panel C and cell lysates were western blotted with the indicated antibodies to determine CS protein disruption. **E.** SLC25A1 depletion is toxic to MYC High cells. MCA data from 4 SLC25A1 sgRNAs and an AAVS1 targeting control. Cells were mixed at a 20%:80% (TRE-MYC:parental) ratio, infected with the SLC25A1 and control sgRNAs shown in panel C, and were examined by flow cytometry after 1 week. **F.** CS depletion is toxic to MYC High cells. MCA data from 4 CS sgRNAs and an AAVS1 targeting control. Cells were mixed as in panel D and cell mixtures were examined by flow cytometry after 1 week. **G.** Colorectal tumor cells with amplified MYC are more sensitive to SLC25A1 disruption by CRISPR. CRISPR screening and copy number data (Absolute) from the Cancer Dependency map (DepMap) were downloaded and analyzed. The p-value was determined by Mann-Whitney test. **H.** MYC-driven increase in lipogenesis requires SLC25A1. TRE-MYC HCECs expressing either sgControl or sgSlc25a1 sgRNAs were treated with vehicle or 100 ng/mL dox for 72 h. LipidTOX green staining was used to determine lipogenesis which was quantified by flow cytometry. Annotation of n.s. denotes non-signifcant, * p=0.0023, ** p=0.0005, and *** p=0.0126 when compared to sgCtl using an unpaired Student’s t-test. Numbers above bar graphs indicate the p-value from an unpaired Student’s t-test to compare sgRNA treated cells to sgControl.

Next we tested whether MYC overexpression can induce synthesis of lipids and whether an increase in lipid synthesis is dependent on SLC25A1 to transport citrate out of the mitochondria. We measured lipid production by cell staining with LipidTOX green which stains neutral lipids and analyzed staining by flow cytometry. In the HCEC TRE-MYC cells, overexpression of MYC led to an increase in LipidTOX staining (Figure 3H). When SLC25A1 was disrupted using CRISPR, the MYC-mediated increase in LipidTOX staining was abrogated indicating a necessity for SLC25A1 for MYC to drive neutral lipid biosynthesis (Figure 3H).

We further confirmed that SLC25A1 and CS are MYC SL genes by employing a second TRE-MYC model using human mammary epithelial cells (HMECs). TRE-MYC HMECs were treated with dox and western blotting showed elevated MYC protein expression (Figure S4A and S4B). Disruption of SLC25A1 and CS expression using CRISPR resulted in a reduction in MYC High cell proliferation as measured by MCA consistent with what was observed earlier in colonic cells (Figure S4C-S4F).

### Inhibition of citrate transport reduces MYC-driven tumor growth in vitro and in vivo

To examine the role of citrate transport in a MYC-driven tumor model, we used M158 mouse mammary tumor cells. M158 cells were derived from a mouse mammary tumor virus (MMTV)-MYC spontaneous mouse breast tumor model (Stewart et al., 1984). We expressed Cas9 and either 2 control sgRNAs targeting mouse intronic sequences or 3 sgRNAs targeting Slc25a1. Western blotting of cell lysates showed a reduction in Slc25a1 expression when Slc25a1 sgRNAs were expressed compared to control (Figure 4A). MCAs comparing the growth of cells with Slc25a1 sgRNAs to cells with control sgRNAs indicated a reduction in growth upon Slc25a1 depletion (Figure 4B). Similar results were obtained when we used sgRNAs targeting mouse citrate synthase, Cs (Figure S5A and S5B). Using shRNAs or sgRNAs targeting mouse Slc25a1 we observed a striking decrease in the anchorage-independent growth of M158 cells in grown in soft agar (Figure S5C-S5E).

**Figure 4.**
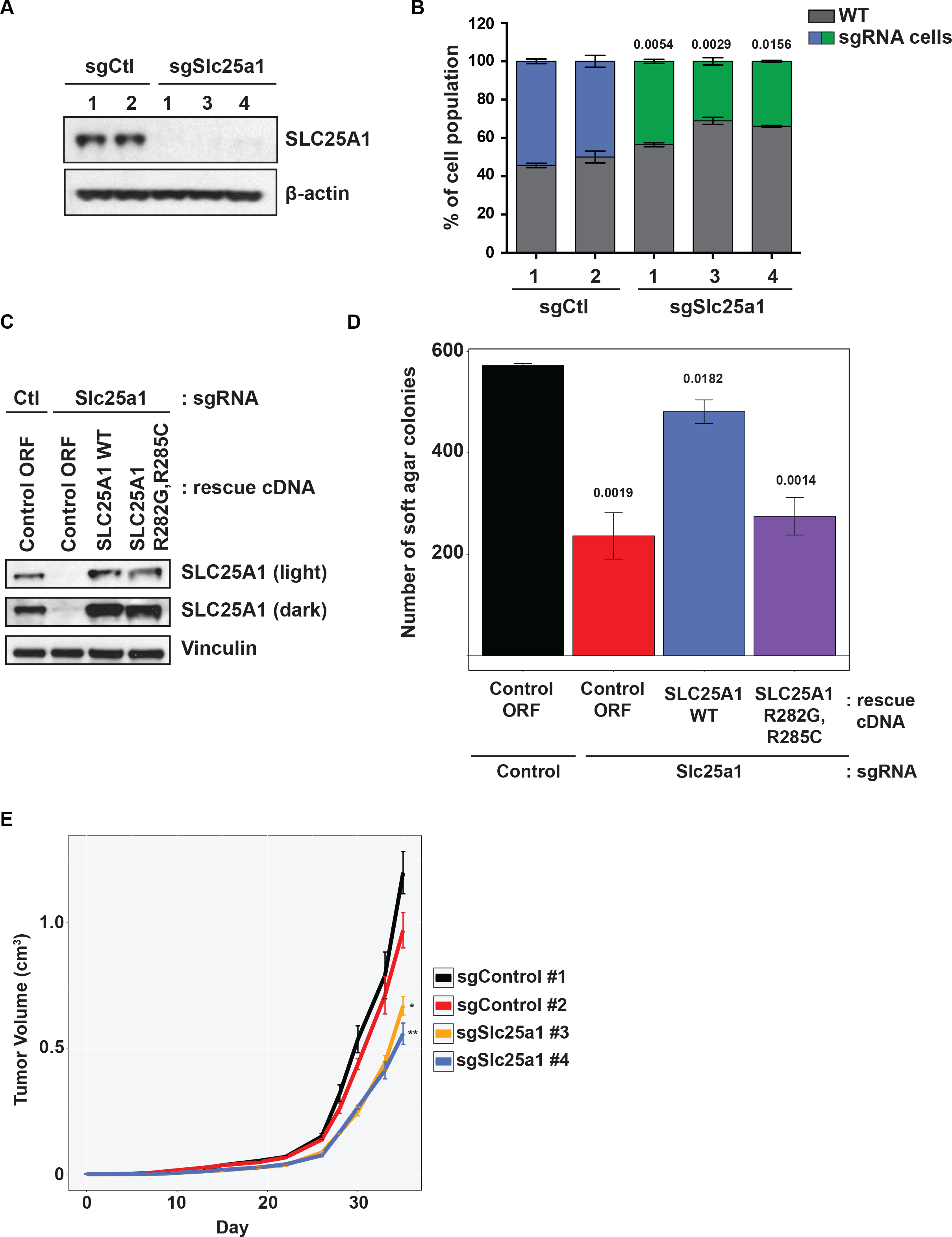
Breast tumor cells driven by MYC overexpression are sensitive to loss of SLC25A1. A. CRISPR targeting of Slc25a1 in M158 cells, a mouse tumor cell line derived from a MMTV-MYC driven breast tumor. M158 cells were infected with Cas9/sgRNA expressing lentiviruses targeting Slc25a1 or safe targeting intron controls. Following 1 week, cell lysates were western blotted with the indicated antibodies to determine Slc25a1 protein disruption. **B.** Loss of Slc25a1 is toxic to M158 cells. GFP+ M158 cells were infected with the indicated sgRNAs. Cells were then mixed with non-fluorescent M158 cells at a 50:50 ratio and allowed to grow for 1 week. Cell populations were determined by flow cytometry. **C.** Rescue of Slc25a1 depletion with WT and citrate binding mutant Slc25a1 cDNAs. M158 cells stably expressing Cas9 and sgSlc25a1 #4 from panel A were infected with lentiviruses to express the indicated cDNAs. Cells were selected with blasticidin and cell lysates were western blotted with the indicated antibodies. The control ORF is the human olfactory receptor OR6F1. **D.** Rescue of the Slc25a1 deficiency induced anchorage independent growth defect. Cells from panel C were plated as single cells in soft agar and colonies were stained with MTT after 14 days. Colonies were counted by imageJ. **E.** Loss of Slc25a1 reduces breast tumor growth. M158 cells expressing the intron targeting control and Slc25a1 #3 and #4 from panel A were subcutaneously implanted into athymic nude female mice. Tumor volume was monitored by caliper measurement every 2 days over the indicated time. * is p=0.0008, ** is p<0.0001 determined from an unpaired Student’s t-test. Numbers above bar graphs indicate the p-value from an unpaired Student’s t-test to compare sgRNA treated cells to sgControl.

Next, we expressed either human SLC25A1 WT or a citrate binding mutant R282G/R285C cDNA in M158 cells where Slc25a1 was depleted by CRISPR (Fernandez et al., 2018). Western blotting showed a similar level of SLC25A1 expression comparable to endogenous Slc25a1 after cDNA expression (Figure 4C). Rescue experiments revealed that the SLC25A1 WT cDNA could restore anchorage-independent growth while the citrate binding mutant failed to do so (Figure 4D).

To examine the effects of targeting Slc25a1 on *in vivo* tumor growth, we subcutaneously implanted M158 cells expressing either intron targeting control or Slc25a1 sgRNAs into athymic nude mice. Tumors that had Slc25a1 sgRNAs showed a reduction in tumor growth indicating a requirement for Slc25a1 in MYC-driven tumor growth (Figure 4E and S5F).

## DISCUSSION

We utilized both CRISPR- and shRNA-based screens to identify the genetic requirements of cells expressing high levels of the oncogene MYC. Core cell growth pathways including nucleotide, lipid, and protein biosynthetic networks all showed preferential toxicity when targeted in cells with elevated MYC (Figure 1E). This is perhaps not surprising given that MYC drives cells into an anabolic state with an increased proliferation rate compared to matched cells without MYC overexpression (Figure 1C). Depletion of nucleotide biosynthesis would be expected to block cellular proliferation but identifying the target enzymes that MYC requires for increased growth without impairing the growth and viability of cells with normal MYC expression provides potential therapeutic targets that may have less toxicity than conventional nucleotide pathway inhibitors. Examples of these types of SL genes from our CRISPR screen include PPAT and ADSS, two components of purine biosynthesis that have not been described as essential genes in previously published CRISPR screens (Hart et al., 2015; Wang et al., 2015). Other MYC SL targets identified in our CRISPR screen include two additional solute carrier (SLC) family transporters like SLC7A5 and SLC33A1 which have been recently described in promoting MYC tumorigenesis (Panda et al., 2020). The requirement for these SLCs is likely due to a necessity to shuttle the metabolic intermediates that MYC uses to drive proliferation. For example, SLC33A1/AT-1 is a transporter that imports acetyl-CoA into the ER where it can be used for protein acetylation (Hirabayashi et al., 2013). SLC33A1 expression is increased when cells have elevated rates of protein synthesis which can cause ER stress and activate the UPR, pathways we found as MYC SL (Figure S1A-C) (Dieterich et al., 2019; Farrugia and Puglielli, 2018). Recently loss of SLC33A1 in KEAP1-deficient lung cancer cells was shown to increase the UPR and increase sensitivity to tunicamycin (Romero et al., 2020). This raises the possibility that loss of SLC33A1 in MYC High cells lethally increases proteotoxic stress creating a targetable vulnerability. SLC33A1 has also been shown to regulate intracellular metabolic crosstalk between the ER and cytosol which alters lipid synthesis indicating that MYC High cells are sensitive to alterations in the balance of acetyl-CoA levels between the ER and cytosol (Dieterich et al., 2019). SLC7A5 is a direct MYC target gene that is upregulated in our RNA-seq data and an amino acid transporter that can activate a feed forward loop to promote MYC expression and tumorigenesis (Figure S1F) (Yue et al., 2017).

Using an alternative to CRISPR based generation of knockouts is necessary to fully investigate SL interactions since CRISPR knockouts of core essential genes is toxic to all cells regardless of genotype. Alternative approaches include generating hypomorphs using CRISPR interference (CRISPRi) or shRNA libraries that target essential genes (Jost et al., 2020). We previously used an shRNA approach to identify essential genes that are SL with oncogenic KRAS (Martin et al., 2017). We generated a new library targeting essential genes using an updated shRNA prediction algorithm (Pelossof et al., 2017) and identified a number of genes that are candidate MYC SL genes. One of these essential SL genes is 6-phosphogluconate dehydrogenase (PGD). By having a deep library with ten shRNAs per gene we are able to have a range of effects due to differing levels of reduction in gene expression. The level of PGD knockdown correlated with the reduction in proliferation (Figures S2C and S2D). This is important since we can define the amount of reduction in these essential genes that is required for cell viability in the presence of excess MYC expression but non-toxic to cells with endogenous MYC levels. PGD encodes for the second dehydrogenase in the pentose phosphate pathway (PPP) which allows cells to generate NAPDH and the ribose necessary for nucleotide biosynthesis (Pavlova and Thompson, 2016). MYC has been shown to facilitate the shuttling of glycolytic intermediates into the PPP (Stine et al., 2015). Competitive Inhibitors of PGD like 6-aminonicotinamide (6-AN) have been examined for cancer therapies as single agents and in combination with conventional chemotherapies like cisplatin (Budihardjo et al., 1998). Acute Myelogenous Leukemia (AML) have frequent MYC amplification and AML cells are sensitive to 6-AN treatment (Chen et al., 2016). Our finding that MYC expression correlates with sensitivity to PPP disruption through inhibition of PGD provides a biomarker for PGD inhibitor sensitivity.

We show that both human and mouse cells with elevated MYC expression derived from breast and colon are reliant upon mitochondrial citrate production and export for growth. Inhibition of citrate synthase or blocking citrate export to the cytosol via SLC25A1 inhibition results in reduced MYC-driven proliferation and tumor growth. A cytosolic pool of acetyl-CoA is essential for the lipogenic needs of cells rapidly proliferating as a result of MYC overexpression. SLC25A1 has been previously described as a MYC interacting protein and a potentiator of cancer cell stemness in non-small cell lung cancer (Fernandez et al., 2018; Koch et al., 2007). Inhibition of a plasma membrane localizing variant of SLC25A1 prevents extracellular uptake of citrate and was shown to reduce fatty acid metabolism and pancreatic tumor growth, a tumor type with elevated expression of MYC that correlates with poor survival (Mycielska et al., 2018). Development of potent and specific SLC25A1 inhibitors may provide a therapy to target tumors driven my high levels of MYC expression.

Acquired resistance to targeted therapies remains a bottleneck to effective tumor therapies and upregulation of MYC is known to drive resistance. For example, MYC protein expression is a biomarker for sensitivity to MAPK inhibition in pancreatic cancer where ERK inhibitor resistance is driven in part through stabilization of the MYC protein (Hayes et al., 2016; Vaseva et al., 2018). Recent work in cells and patient-derived xenografts (PDX) with acquired resistance to bromodomain inhibition by JQ-1 through MYC upregulation has shown that inhibition of anti-apoptotic proteins including MCL1, one of our identified MYC SL genes (Figure 2C and 2D), results in hypersensitivity and reduced tumor growth (Lin et al., 2020). It is possible that combinations of the inhibitors of the SL genes we have identified with either MAPK or bromodomain inhibitors will prevent MYC-dependent acquired resistance in tumor cells or enhance their lethality at lower concentrations.

From our CRISPR screens, we also observed that a significant number of genes when mutant allow MYC High cells to proliferate even better relative to WT. Loss of function mutations of these genes are synthetic enhancers of proliferation in MYC High cells and one would predict that this class of genes is more frequently mutated in MYC amplified tumors. One of these genes is *NFE2L2* which encodes the redox regulator protein NRF2. Our CRISPR screens found that disruption of NRF2 expression is toxic in MYC WT cells while providing a growth advantage for MYC High cells. Previous work in mouse tumor models has shown that loss of NRF2 enhances the tumorigenicity of APC(min/+) mice and *NFE2L2* is more frequently mutated in MYC amplified tumors (Cheung et al., 2014). High levels of proliferation due to increased MYC expression generate reactive oxygen species (ROS) that can reduce cell growth. Loss of the NRF2 redox sensor in MYC High cells may enable bypass ROS sensing helping to further increase MYC-driven proliferation. While these synthetic enhancer genes aren’t therapeutic targets to block MYC induced growth, the performance of genes like *NFE2L2* underscore how increased MYC expression and rapid proliferation alters cellular stress networks.

MYC, as a master regulator of cell growth, must initiate the coordination of multiple cellular pathways that generate the material required to duplicate a cell. This coordination implies a balance that must be established so that different aspects of the process move forward together. Reducing only a portion of the program can cause an imbalance that could derail the entire program and lead to toxic outcomes. Inhibition of downstream anabolic pathways including lipid synthesis, nucleotide metabolism, splicing, or protein translation all selectively penalize MYC overproducing cells that are attempting to force bulk metabolic activation to maximize cell proliferation. It is clear that the MYC High cells, while growing fast, are not doing so completely optimally and by forcing down the cell proliferation accelerator pedal, so to speak, are stressing the fidelity of some processes. DNA replication is likely to be one of those given the identification of DBF4 and MCM10 as MYC SL genes. For this reason, MYC High cells are reliant on stress support pathways such as DNA repair, the DNA damage response pathway, and the UPR. This requirement for stress response networks creates therapeutically targetable NOA whose inhibition can selectively block MYC High cell proliferation. As described in Luo et al. (2009), cases of

NOA that depend on stress response pathways can be targeted by either creating additional stress so as to overwhelm the stress response pathway as we observed with the UPR inducer tunicamycin (Figure S1C) or by reducing the functionality of the stress response pathway as we showed with Rad17 for DNA replication and HSF1 for protein folding. Again, these agents provide an orthogonal therapeutic approach that could be combined with other therapeutics to enhance lethality in MYC high tumors. Future studies are needed to examine the utility of inhibitor combinations and dosing regimens that concurrently inhibit both the MYC SL targets like SLC25A1 along with NOA networks in combatting MYC-driven malignancies.

## EXPERIMENTAL PROCEDURES

### Cell Culture and Reagents

Human colonic epithelial cells (HCECs) were obtained from Dr. Jerry Shay (UT- Southwestern) and were cultured as described (Roig et al., 2010). HEK293Ts were obtained from ATCC (CRL-3216), grown in DMEM with 10% fetal calf serum (HyClone) and 1% penicillin/streptomycin (Gibco), and were used to generate lentiviruses. M158 mouse mammary tumor cells were obtained from ATCC (CRL-3086), grown in DMEM with 10% fetal calf serum (HyClone) and 1% penicillin/streptomycin (Gibco). Human mammary epithelial cells (HMECs) have been previously described (Sack et al., 2018). All cells were tested for Mycoplasma prior to and routinely during use (Lonza, cat. # LT07- 218).

### Plasmids and other reagents

Human MYC and SLC25A1 cDNAs were from the Ultimate ORF collection (Thermo) and were in pDONR221. MYC and SLC25A1 cDNAs were recombined in LR reactions into a pInducer20 (Meerbrey et al., 2011) variant with blasticidin selection. Expression of cDNAs was induced with 100 ng/mL doxycycline (Clontech/Takara). Site-directed mutagenesis was used to generate the citrate binding mutant SLC25A1. sgRNAs were ordered as complimentary oligos with 5’-CACC overhangs on the forward strand and 5’-AAAC overhangs on the reverse strand. Annealed and phosphorylated oligos were ligated into lentiCRISPR V2 puro with an FE modified tracr (Chen et al., 2013).

shRNAs were ordered as 97-mers and PCR amplified with mir-E sequences and XhoI/EcoI restriction sites. Digested PCR products were ligated into a doxycycline-inducible pHAGE ind10 mirE lentiviral backbone (Martin et al., 2017). Expression of shRNAs was induced with 100 ng/mL dox. Sequences of the sgRNAs and shRNAs used are described in Supplemental Table 5.

### Genome-Wide CRISPR Screening For MYC Synthetic Lethal Genes

A library of gRNAs targeting 18,166 genes with 5 gRNAs per gene for a total of approximately 91,000 gRNAs was previously described (Martin et al., 2017). Pooled virus was prepared by transfecting 293T cells with the library plasmid pool and psPax2 and pMD2.G lentiviral packaging vectors. Viral supernatants were harvested at 48 and 72 hours post transfection and concentrated with lenti-X concentrator solution (Clontech). TRE-MYC HCECs were infected in triplicate at a low MOI (0.2) with a representation of 500 cells per sgRNA. Cells were selected with puromycin (1 μg/mL) for 3 days until an uninfected control plate completely died. An initial cell pellet was taken as PD0, cells were split into no dox and 100 ng/mL dox treated groups, and cells were grown an additional 5 and 10 population doublings (PD5 and PD10), where a sample of cells was collected. Genomic DNA was isolated by phenol:chloroform extraction and gRNAs were PCR amplified with barcoded primers for sequencing on an Illumina NextSeq 500. Sequencing reads were aligned to the CRISPR library and raw read counts were obtained for each sgRNA. MAGeCK and edgeR were used to normalize reads, calculate p-values, FDRs, and log2 fold changes for comparison between the PD5 and PD10 with the input PD0 samples. GSEA analyses were performed using the Java based GSEA tool (version 4.0.4) and MSigDB curated gene sets (The Broad Institute).

### shRNA Essential Gene Screen

A library of ∼12,000 shRNAs targeting 1,111 essential genes with ∼10 shRNAs per gene derived from the SPLASH shRNA dataset (Pelossof et al., 2017) was synthesized via oligo array and cloned into a doxycycline-inducible pHAGE ind10 mirE vector. One thousand shRNAs targeting *E. coli* genes were included as negative controls. Virus was prepared as for the CRISPR screen and the HCECs were infected at a MOI of 0.2 in triplicate with a representation of 1,000 cells per sgRNA. Following puromyin selection, an initial PD0 cell pellet was taken and cells were treated with 100 ng/mL doxycycline to induce shRNA expression. Cells were maintained in 100 ng/mL doxycycline throughout the screen and cell pellets were taken at PD3, PD7, and PD10 to monitor the abundance of shRNAs over time. Genomic DNA isolation, sequencing of shRNAs, and screen data analyses were performed as described for the CRISPR screen.

### LipidTOX green staining

One hundred thousand cells were plated in 12-well plates in triplicate and either treated with DMSO or 100 ng/mL doxycycline to induce MYC expression. After 72h of MYC induction, live cells were incubated with LipidTOX HCS Green (ThermoFisher, cat# H34475) for 30 min at a 1:500 dilution. Cells were washed with PBS, trypsinized and placed on ice, and examined for lipid staining by flow cytometry using a LSRII (BD) flow cytometer. The geometric mean of the LipidTOX green signal was used to quantify lipid accumulation.

### Proliferation and Soft Agar Assays

For multicolor competition assays (MCAs), fluorescent and non-fluorescent cells were mixed at the indicated ratios and examined by flow cytometry on a LSRII (BD) directly after mixing and after the indicated time points described in the figures. Green fluorescent HCECs were generated by infecting with pHAGE blast CMV mClover3 PGK blast lentiviruses and sorting the fluorescent population. Green fluorescent M158 cells were generated by infecting with pHAGE EF1 mClover3 PGK blast and sorting the fluorescent population.

1,000 HCECs or M158 cells were plated per well of 96-well plates and proliferation was measured over time using CellTiter Blue (Promega) and a plate reader. GR50 values for the indicated drugs were determined using the online GR calculator (http://www.grcalculator.org/grcalculator/) (Hafner et al., 2016).

For soft agar assays, 10,000 cells in a layer of 0.6% agar were plated on a bottom layer of 0.6% agar and allowed to grow for 2-3 weeks. Colonies were stained with MTT (5 mg/mL, Sigma) and quantified using imageJ (NIH).

### Western blotting and antibodies

All cells were lysed in 1x RIPA lysis buffer (50 mM Tris-HCl, 150 mM NaCl, 1% NP-40, 0.5% sodium deoxycholate and 0.1% SDS, Boston Bioproducts, cat. # BP-115X) with protease and phosphatase inhibitors (Thermo, cat. # 78440). Protein was quantified by Bradford assay (Bio-Rad, cat. # 500-0006EDU). Protein lysates were mixed with 4X reducing sample buffer (Invitrogen, cat. # NP0007), separated on 4-12% Bis-Tris gels (Thermo, cat. # NP0336BOX), and transferred via a Trans-Blot Turbo Transfer System (Bio-Rad, cat. # 1704150) to nitrocellulose membranes (Bio-Rad, cat.# 170-4158). Membranes were blocked for 1 h at room temperature with 5% milk in TBST (Santa Cruz, cat. # sc-362311) and incubated with the following antibodies at the indicated dilutions in 5% BSA (VWR, cat. # VWRV0332) in TBST either overnight at 4C or 1 h at room temperature: Vinculin (1:10,000, Sigma cat. # V9131), MYC (1:1,000, Cell Signaling, cat. # 5605), CHKA (1:1,000, Cell Signaling, cat. # 13422), CS (1:1,000, ProteinTech, cat. # 16131-1-AP), SLC25A1 (1:1,000, ProteinTech, cat. # 5235-1-AP), PGD (1:1,000, ProteinTech, cat. # 14718-1-AP), ATF4 (1:1,000, Cell Signaling, cat. # 11815), CHOP (1:1,000, Cell Signaling, cat. # 2895), PARP (1:1,000, Cell Signaling, cat. # 9542), Caspase 3 (1:1,000, Cell Signaling, cat. # 9662), HSF1 (1:1,000, Cell Signaling, cat. # 4356), β-actin(1:10,000, Cell Signaling, cat. # 3700), and GAPDH (1:10,000, Santa Cruz, cat. # sc-365062).

Goat anti-mouse or rabbit HRP conjugated secondary antibodies (1:5,000, Fisher, cat. #s 31430 and 31460) incubated for 1 h at room temperature were used for detection with ECL (Perkin-Elmer, cat. # NEL104001EA) via LI-COR (Odyssey model #2800) or film (HyBlot CL, Denville Scientific, cat. # 1159M38).

### Animal Xenografts Assays

For all xenografts, 1 million M158 cells in 100uL PBS (Gibco) were injected subcutaneously into 8-week old female athymic mice (Stock # 002019, The Jackson Laboratory). Tumors were measured by caliper every 2 days. Tumor volume was calculated by ((length) x (width) x (width))/2. All animal studies were done with an IACUC approved protocol.

### Statistical Analyses

Differences between two groups were determined using an unpaired, two-tailed Student’s t test in Prism (Graphpad) unless otherwise noted. Error bars denote S.E.M. All assays were performed in duplicate or triplicate as indicated and repeated for a total of at least two independent experiments.

## AUTHOR CONTRIBUTIONS

T.D.M. and M.Y.C performed genome-wide CRISPR and essential gene shRNA screens, western blotting, cell viability and growth assays, and shRNA experiments; T.D.M. analyzed the screening data; T.D.M. and M.Z.L. prepared the shRNA and CRISPR lentiviral libraries; T.D.M., R.P. and A.L. performed all *in vivo* experiments; T.D.M. and S.J.E. conceived the study and wrote the paper with input from all authors.

## ACKNOWLEDGMENTS

T.D.M. was supported by a Damon Runyon Cancer Research Foundation fellowship (DRC 2199-14). We thank Eric Wooten for help with bioinformatic analysis and members of the Elledge lab for helpful discussions. This work was supported by a grant from the NCI and the Ludwig Foundation to S.J.E. S.J.E. is an Investigator with the Howard Hughes Medical Institute.

---

**Supplemental Figure 1.**
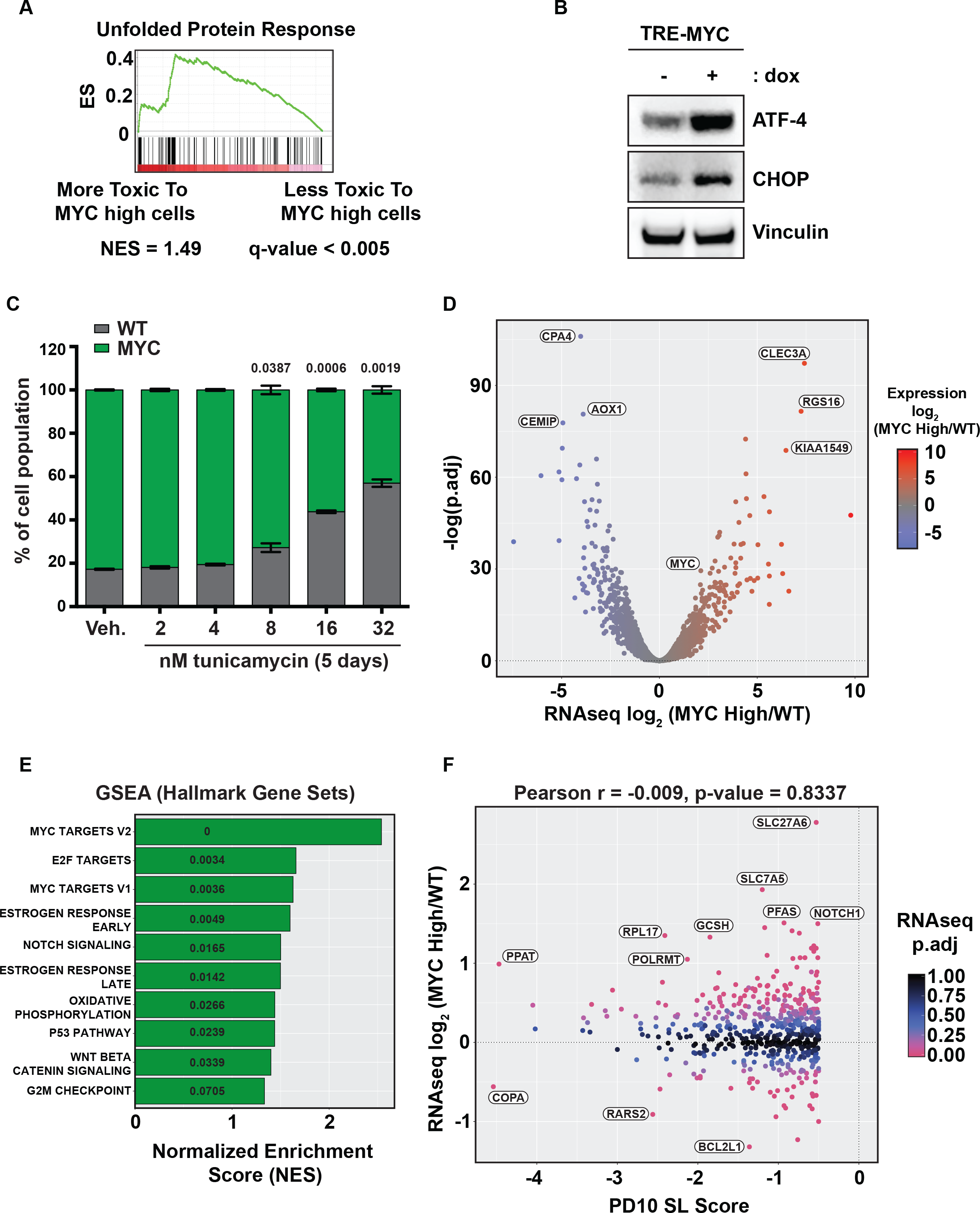
Further characterization of the MYC synthetic lethal CRISPR screen. **A.** Disruption of the unfolded protein response (UPR) is more toxic to MYC overexpressing cells. GSEA plot of genes in the Hallmark gene set related to the UPR. **B.** Overexpression of MYC increases the expression ATF4 and CHOP, two components of the UPR and ER stress pathways. TRE-MYC HCECs were treated with vehicle or 100 ng/mL doxycycline for 72 h. Cell lysates were western blotted for the indicated proteins. **C.** Cells with high expression of MYC are more sensitive to the UPR and ER stress inducing drug tunicamycin. Non-fluorescent TRE-MYC cells were mixed with GFP expressing parental cells at a 20%:80% (TRE-MYC:parental) ratio and treated with the indicated doses of tunicamycin. After 5 days cell populations were examined by flow cytometry. **D.** Transcriptome analysis of the TRE-MYC cells used in the CRISPR screen. TRE-MYC HCEC cells used in the screen were treated with 100 ng/mL doxycycline for 72 h. Total mRNA was isolated and used for cDNA library preparation and RNA-seq. The volcano plot includes some of the genes whose expression is most increased (including MYC itself) and decreased as a result of MYC overexpression. Full RNA-seq data are available as Supplemental Table 3. **E.** GSEA of pathways upregulated after MYC overexpression. RNA-seq data were used for GSEA to identify pathways of genes that are increased after MYC induction. **F.** No correlation of the MYC transcriptome with the results from the MYC synthetic lethal screen. Data from the TRE-MYC RNA-seq were compared to the synthetic lethal CRISPR screening results where the PD10 SL Score is the log2 fold change difference between MYC High and MYC WT cells at PD10 (High-WT). Pearson correlation coefficient and p-values were determined.

**Supplemental Figure 2.**
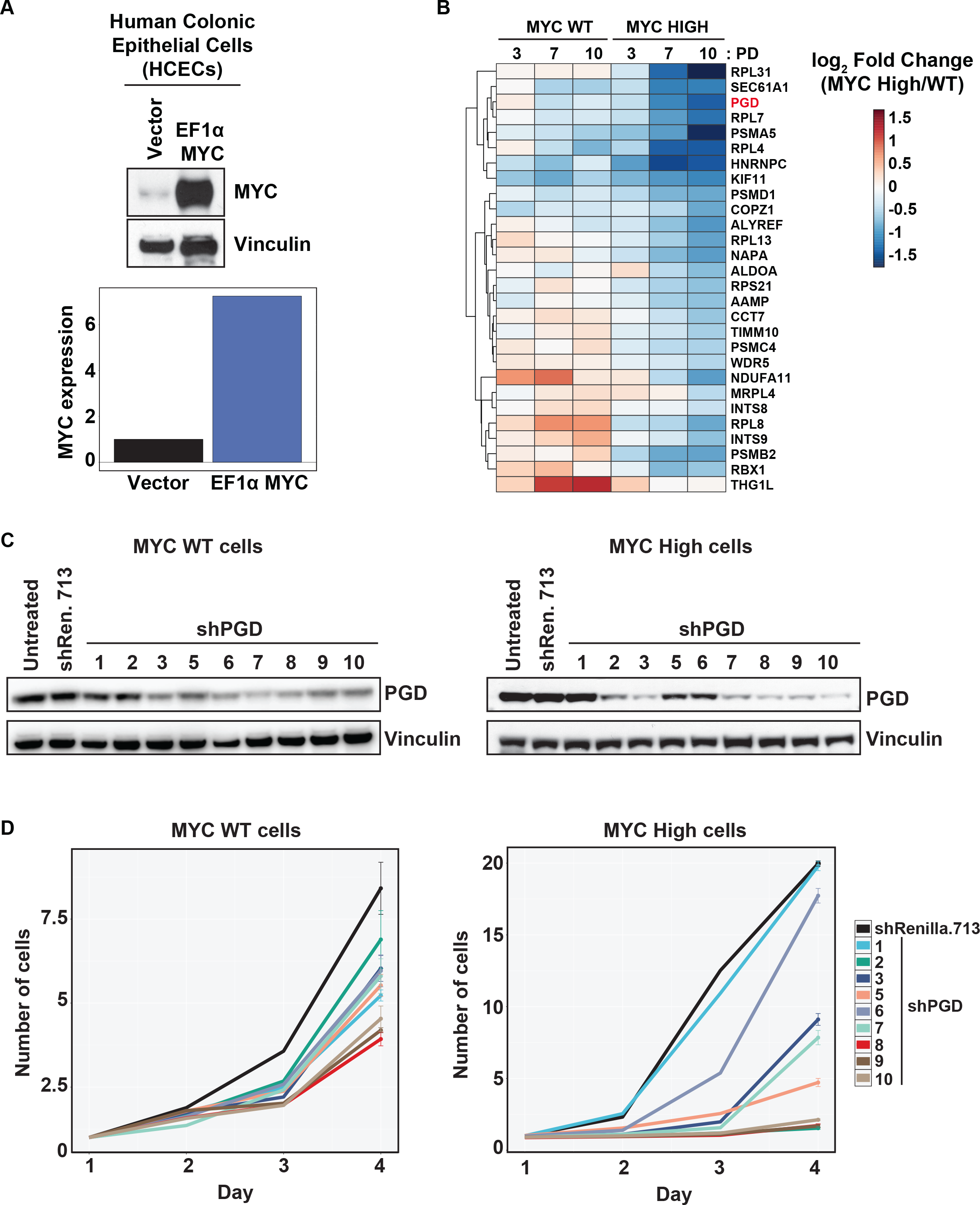
An essential gene shRNA screen to identify MYC SL genes missed by CRISPR-based approaches. **A.** Generation of HCECs with constitutive expression of MYC. HCECs were infected with EF1-MYC PGK blast lentiviruses and selected with blasticidin. Cell lysates were examined for MYC expression by western blotting with the indicated antibodies and quantified using LI-COR. **B.** Top scoring essential genes that are differentially required for MYC overexpressing cells. Cells from panel A were infected with a lentiviral dox inducible shRNA library targeting essential genes which are universally toxic in CRISPR screens. Cells were sequenced at PD 3, 7, and 10 to monitor shRNA abundance. This heatmap shows the top scoring genes that are more required in the context of MYC overexpression. **C.** shRNA-mediated knockdown of 6-phosphogluconate dehydrogenase (PGD) expression. Cells from panel A were infected with dox inducible shRNAs targeting PGD or luciferase (shRen.713) control. Cells were treated with 100 ng/mL doxycycline for 72 h and cell lysates were western blotted with the indicated antibodies to measure PGD expression. **D.** PGD is an essential MYC synthetic lethal gene. MYC WT and MYC High HCECs expressing the indicated shRNAs were plated in 96-well plates, treated with 100 ng/mL doxycycline to induce shRNA expression, and cell proliferation was monitored by CellTiter Blue over 4 days.

**Supplemental Figure 3.**
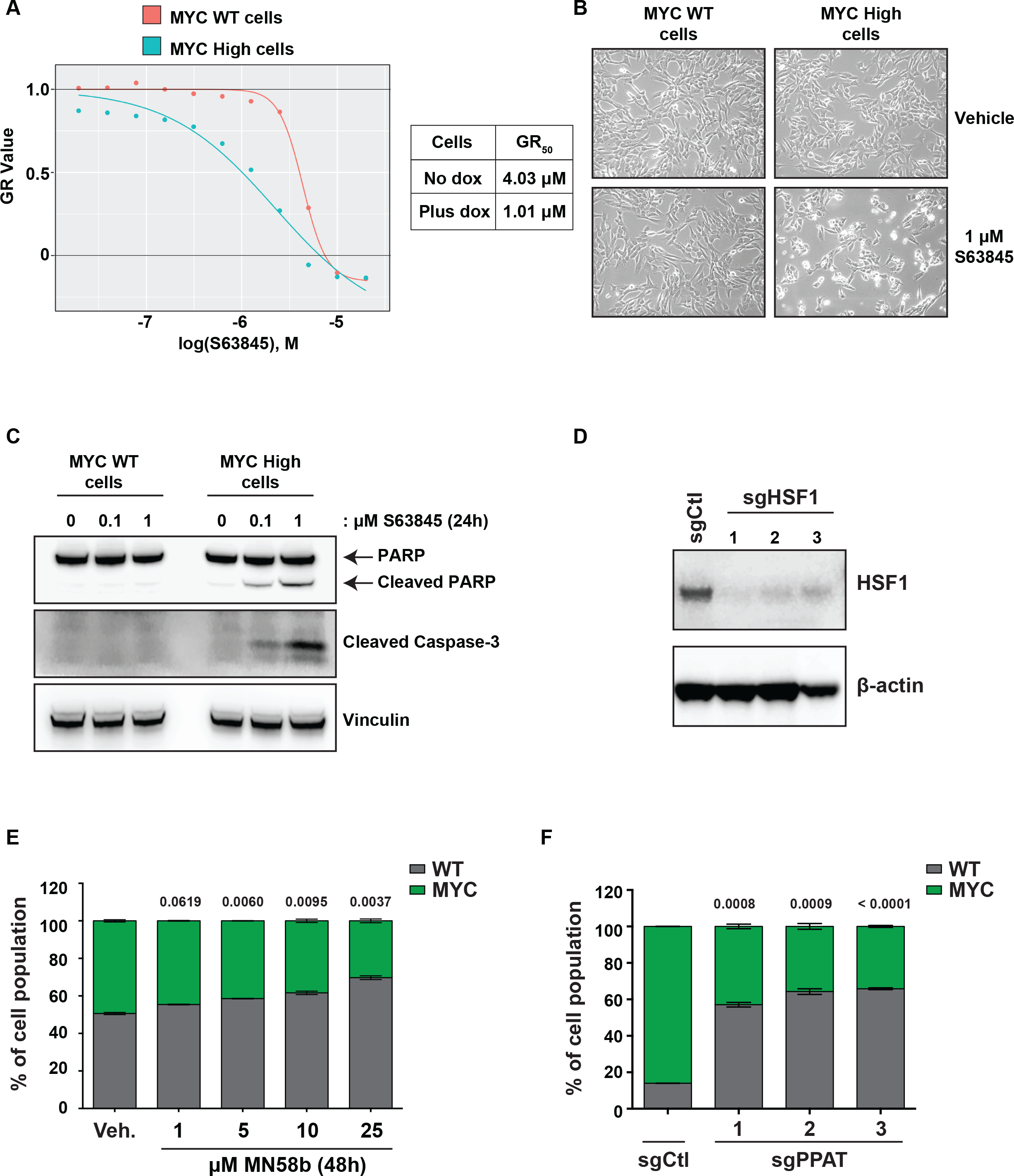
Further validation of a selection of MYC SL genes. **A.** MYC overexpression increases sensitivity to the MCL1 inhibitor S63845. HCEC cells with TRE-MYC were treated vehicle or 100 ng/mL doxycycline and with the indicated doses of S63845. After 72 h, cell viability was measured by CellTiter Blue and the GR50 was determined using the online GR calculator (http://www.grcalculator.org/grcalculator/) which can correct for the proliferation rate difference in cells with low and high MYC expression. **B.** Treatment of MYC overexpressing cells with S63845 induces visual apoptosis. Cells were treated with 1 μM of S63845 for 72 h and images were taken. Cells treated with dox to induce high levels of MYC expression were noticeably more apoptotic (i.e. round cells that detached from the plate) than those not treated with dox. **C.** Treatment of MYC overexpressing cells with S63845 induces apoptosis as determined by western blot. HCEC cells with TRE-MYC were treated vehicle or 100 ng/mL doxycycline and either DMSO, 100 nM, or 1 μM S63845 for 24 h. Cell lysates were analyzed for cleaved caspase 3 and cleaved PARP to monitor apoptosis induction. **D.** CRISPR disruption of HSF1 expression. HSF1 targeting sgRNAs used in the MCA in Figure 2E were expressed in HCEC TRE-MYC cells. Following 1 week of sgRNA expression, HSF1 protein levels were determined by western blot. **E.** Inhibition of CHKA via MN58b is more toxic to MYC overexpressing cells. Non-fluorescent HCEC TRE-MYC cells were mixed with GFP+ HCECs at an 80:20 ratio. Cell mixtures were treated with the indicated concentrations of the CHKA inhibitor MN58b or DMSO vehicle control. Changes in cell populations were determined after 48 h of treatment by flow cytometry. **F.** CRISPR disruption of PPAT is toxic to MYC overexpressing cells. Non-fluorescent HCEC TRE-MYC cells were mixed with GFP+ HCECs at an 80:20 ratio. Cell mixtures were treated with the indicated sgRNAs. After 1 week in 100 ng/mL dox, cell populations were examined by flow cytometry. Numbers above bar graphs indicate the p-value from an unpaired Student’s t-test to compare sgRNA or treatment groups to sgControl or Vehicle, respectively.

**Supplemental Figure 4.**
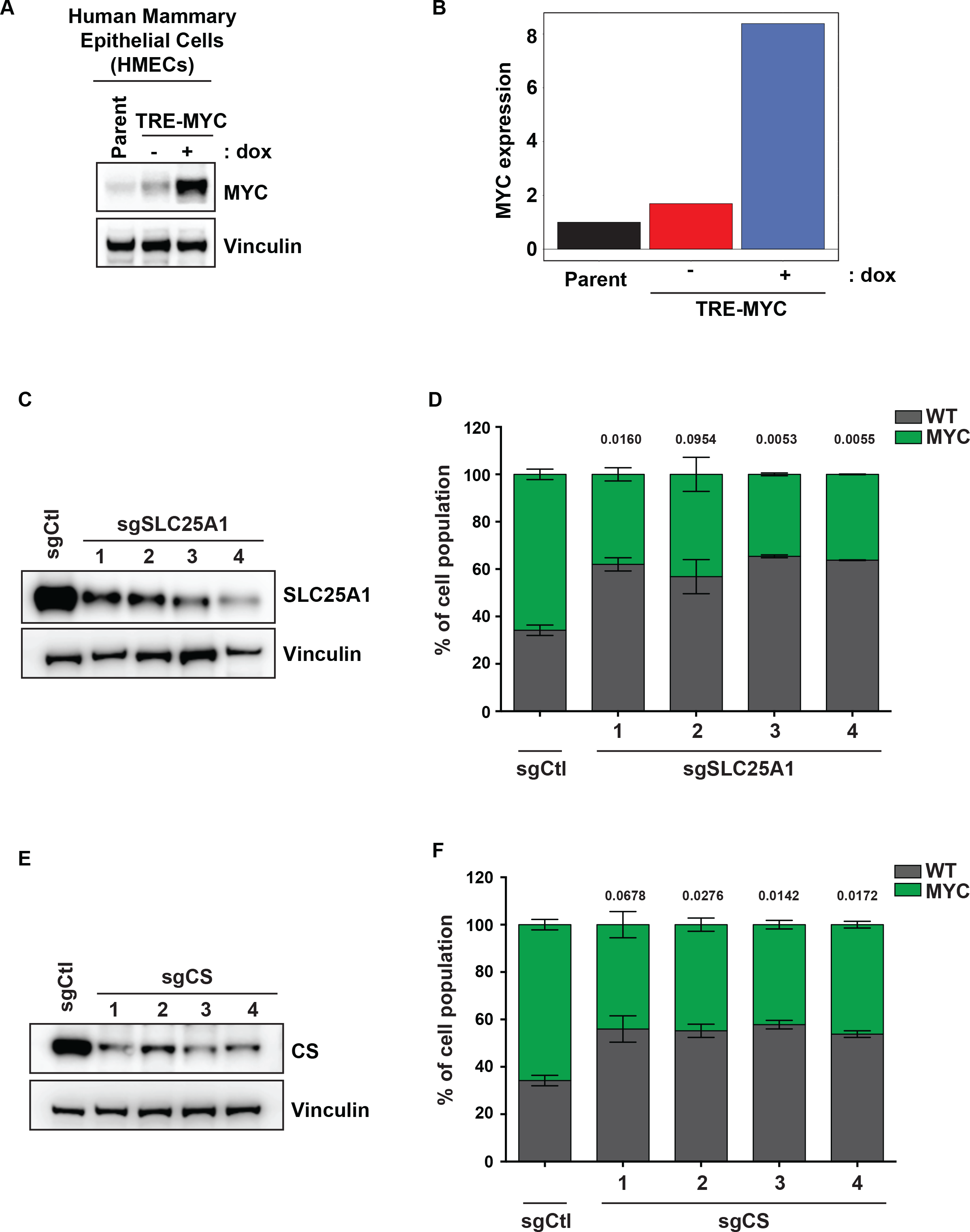
Validation of MYC SL genes in human mammary epithelial cells. **A.** Generation of TRE-MYC human mammary epithelial cells (HMECs). HMECs were infected with a pInducer20-based lentivirus containing TRE-MYC UbC-rtTA3-IRES-blast to generate the TRE-MYC cell line. Blasticidin resistant cells were examined for MYC expression after 48 h of 100 ng/mL doxycycline treatment. **B.** Quantification of MYC induction. Bar graph representing the fold induction of MYC before and after dox treatment as determined by LI-COR quantification of the western blot in panel A. **C.** CRISPR disruption of SLC25A1 in TRE-MYC HMECs. TRE-MYC HMECs were infected with the indicated sgRNA constructs. After 7 days, cell lysates were western blotted to determine loss of SLC25A1 expression. **D.** Loss of SLC25A1 is toxic to MYC overexpressing HMECs. Non-fluorescent TRE-MYC HMECs were mixed with GFP+ parental HMECs at a 20:80 ratio. Cell populations were infected with the indicated sgRNAs, treated with 100 ng/mL dox, and mixtures were examined by flow cytometry 7 days later. **E.** CRISPR disruption of CS in TRE-MYC HMECs. TRE-MYC HMECs were infected with the indicated sgRNA constructs. After 7 days, cell lysates were western blotted to determine loss of CS expression. **F.** Loss of CS is toxic to MYC overexpressing HMECs. Cells were mixed as in panel D, infected with the indicated sgRNAs, treated with 100 ng/mL dox, and cell mixtures were examined by flow cytometry 7 days later. Numbers above bar graphs indicate the p-value from an unpaired Student’s t-test to compare sgRNA treated cells to sgControl.

**Supplemental Figure 5.**
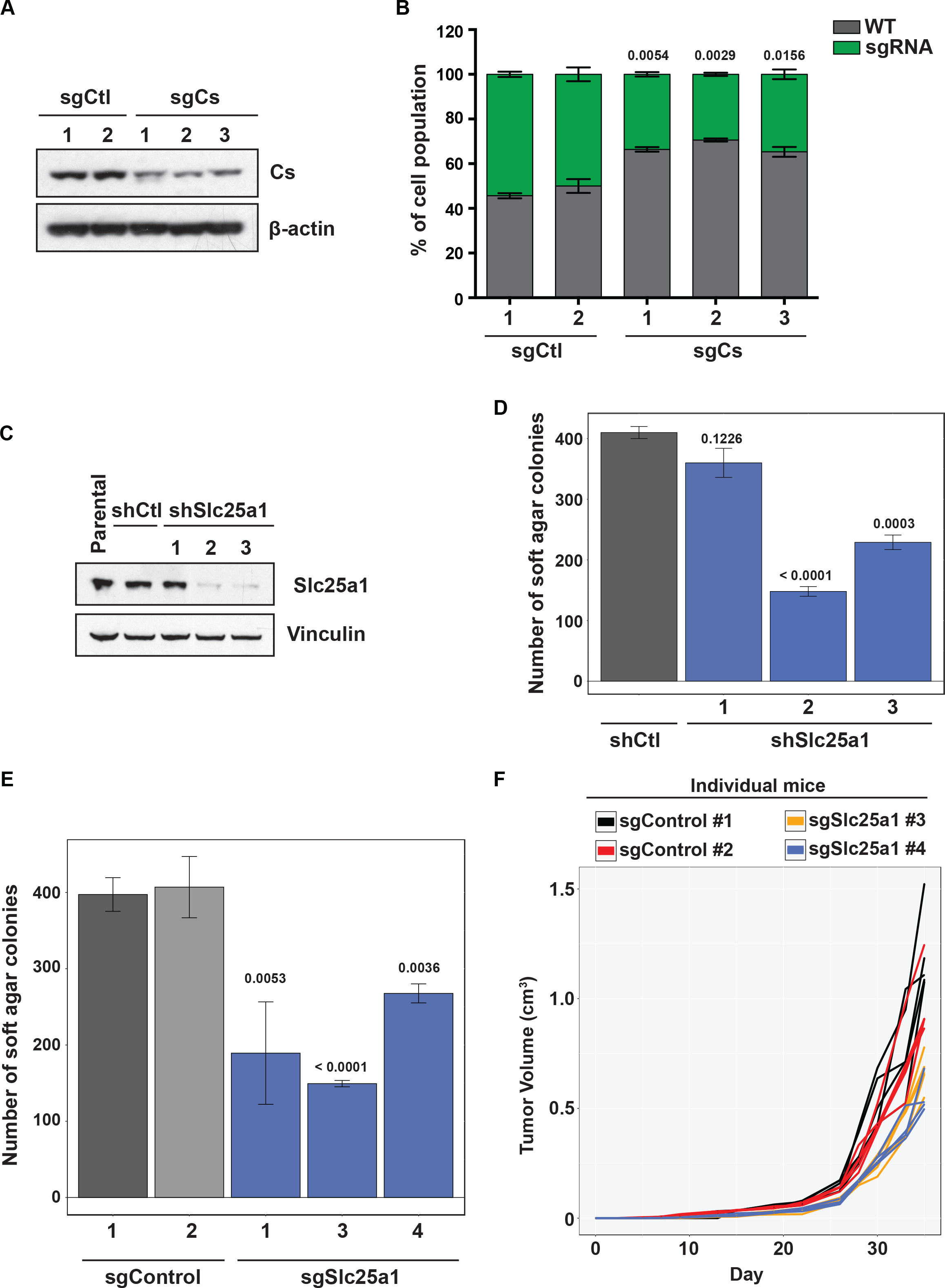
G**e**netic **perturbation of mitochondrial citrate production and transport reduces MYC-driven breast tumor cell growth.** A. CRISPR-mediated disruption of citrate synthase. MYC-driven M158 mouse mammary tumor cells were infected with the indicated sgRNAs targeting Cs or mouse intron targeting cutting controls (sgCtl). One week after infection, cell lysates were western blotted to determine loss of Cs expression. B. Loss of Cs is toxic to M158 breast tumor cells. GFP+ M158 cells were infected with the indicated sgRNAs. Cells were then mixed with non-fluorescent M158 cells at a 50:50 ratio and allowed to grow for 1 week. Cell populations were determined by flow cytometry. C. RNAi targeting Slc25a1 transcripts reduces growth of M158 cells. Dox inducible shRNA hairpins including three targeting Slc25a1 or a control targeting luciferase (Renilla 713) were expressed in M158 cells. Cells were treated with 100 ng/mL doxycycline for 72 h and cell lysates were western blotted to determine loss of Slc25a1 protein expression. D. Loss of Slc25a1 expression by RNAi reduces M158 soft agar growth. Cells from panel C were placed in soft agar and colony formation was examined 2 weeks later by MTT staining and quantification using imageJ. E. Loss of Slc25a1 expression by CRISPR reduces M158 soft agar growth. M158 cells expressing the indicated sgRNAs were placed in soft agar and colony formation was examined 2 weeks later as in panel D. F. Slc25a1 targeting by CRISPR reduces M158 tumor growth in vivo. M158 cells expressing intron targeting control sgRNAs or sgRNAs targeting Slc25a1 (#3 and #4) were subcutaneously implanted into athymic nude female mice with 5 mice per group. Tumor volume was monitored by caliper measurement every 2 days over the indicated time. Data are the individual mouse tumor volumes used to generate the graph in Figure 4F. |Numbers above bar graphs indicate the p-value from an unpaired Student’s t-test to compare shRNA or sgRNA treated cells to shControl or sgControl, respectively.

**Supplemental Table 1. Raw read counts for genome-wide CRISPR screen –** Raw aligned read counts for the HCEC TRE-MYC CRISPR SL screen used for edgeR and MAGeCK analyses

**Supplemental Table 2. CRISPR screen analysis of TRE-MYC HCECs –** edgeR and MAGeCK analyses of the HCEC TRE-MYC CRISPR SL screen including a list of putative SL genes

**Supplemental Table 3. RNA-seq analysis of TRE-MYC HCECs –** Raw aligned read counts for HCEC TRE-MYC RNA-seq and analyses including GSEA

**Supplemental Table 4. shEssential screen analysis of TRE-MYC HCECs –** Raw aligned read counts for the HCEC EF1-MYC shEssential screen and edgeR analyses

**Supplemental Table 5. Sequences of sgRNAs and shRNAs used for validation experiments –** Sequences of sgRNAs and shRNAs used throughout the figures

## REFERENCES

1. Beaulieu, M.E., Jauset, T., Masso-Valles, D., Martinez-Martin, S., Rahl, P., Maltais, L., Zacarias-Fluck, M.F., Casacuberta-Serra, S., Serrano Del Pozo, E., Fiore, C., et al. (2019). Intrinsic cell-penetrating activity propels Omomyc from proof of concept to viable anti-MYC therapy. Sci Transl Med 11.

2. Bryant, H.E., Schultz, N., Thomas, H.D., Parker, K.M., Flower, D., Lopez, E., Kyle, S., Meuth, M., Curtin, N.J., and Helleday, T. (2005). Specific killing of BRCA2-deficient tumours with inhibitors of poly(ADP-ribose) polymerase. Nature 434, 913–917.

3. Budihardjo, II, Walker, D.L., Svingen, P.A., Buckwalter, C.A., Desnoyers, S., Eckdahl, S., Shah, G.M., Poirier, G.G., Reid, J.M., Ames, M.M., et al. (1998). 6-Aminonicotinamide sensitizes human tumor cell lines to cisplatin. Clin Cancer Res 4, 117–130.

4. Cermelli, S., Jang, I.S., Bernard, B., and Grandori, C. (2014). Synthetic lethal screens as a means to understand and treat MYC-driven cancers. Cold Spring Harb Perspect Med 4.

5. Chen, B., Gilbert, L.A., Cimini, B.A., Schnitzbauer, J., Zhang, W., Li, G.W., Park, J., Blackburn, E.H., Weissman, J.S., Qi, L.S., et al. (2013). Dynamic imaging of genomic loci in living human cells by an optimized CRISPR/Cas system. Cell 155, 1479–1491.

6. Chen, Y., Xu, Q., Ji, D., Wei, Y., Chen, H., Li, T., Wan, B., Yuan, L., Huang, R., and Chen, A.(2016). Inhibition of pentose phosphate pathway suppresses acute myelogenous leukemia. Tumour Biol 37, 6027–6034.

7. Cheung, K.L., Lee, J.H., Khor, T.O., Wu, T.Y., Li, G.X., Chan, J., Yang, C.S., and Kong, A.N. (2014). Nrf2 knockout enhances intestinal tumorigenesis in Apc(min/+) mice due to attenuation of anti-oxidative stress pathway while potentiates inflammation. Mol Carcinog 53, 77–84.

8. Dai, C., Whitesell, L., Rogers, A.B., and Lindquist, S. (2007). Heat shock factor 1 is a powerful multifaceted modifier of carcinogenesis. Cell 130, 1005–1018.

9. Dang, C.V., Le, A., and Gao, P. (2009). MYC-induced cancer cell energy metabolism and therapeutic opportunities. Clin Cancer Res 15, 6479–6483.

10. Dieterich, I.A., Lawton, A.J., Peng, Y., Yu, Q., Rhoads, T.W., Overmyer, K.A., Cui, Y., Armstrong, E.A., Howell, P.R., Burhans, M.S., et al. (2019). Acetyl-CoA flux regulates the proteome and acetyl-proteome to maintain intracellular metabolic crosstalk. Nat Commun 10, 3929.

11. Farmer, H., McCabe, N., Lord, C.J., Tutt, A.N., Johnson, D.A., Richardson, T.B., Santarosa, M., Dillon, K.J., Hickson, I., Knights, C., et al. (2005). Targeting the DNA repair defect in BRCA mutant cells as a therapeutic strategy. Nature 434, 917–921.

12. Farrugia, M.A., and Puglielli, L. (2018). Nepsilon-lysine acetylation in the endoplasmic reticulum - a novel cellular mechanism that regulates proteostasis and autophagy. J Cell Sci 131.

13. Felsher, D.W., and Bishop, J.M. (1999). Reversible tumorigenesis by MYC in hematopoietic lineages. Mol Cell 4, 199–207.

14. Fernandez, H.R., Gadre, S.M., Tan, M., Graham, G.T., Mosaoa, R., Ongkeko, M.S., Kim, K.A., Riggins, R.B., Parasido, E., Petrini, I., et al. (2018). The mitochondrial citrate carrier, SLC25A1, drives stemness and therapy resistance in non-small cell lung cancer. Cell Death Differ 25, 1239–1258.

15. Gad, H., Koolmeister, T., Jemth, A.S., Eshtad, S., Jacques, S.A., Strom, C.E., Svensson, L.M., Schultz, N., Lundback, T., Einarsdottir, B.O., et al. (2014). MTH1 inhibition eradicates cancer by preventing sanitation of the dNTP pool. Nature 508, 215–221.

16. Gao, P., Tchernyshyov, I., Chang, T.C., Lee, Y.S., Kita, K., Ochi, T., Zeller, K.I., De Marzo, A.M., Van Eyk, J.E., Mendell, J.T., et al. (2009). c-Myc suppression of miR-23a/b enhances mitochondrial glutaminase expression and glutamine metabolism. Nature 458, 762–765.

17. Gutteridge, R.E., Ndiaye, M.A., Liu, X., and Ahmad, N. (2016). Plk1 Inhibitors in Cancer Therapy: From Laboratory to Clinics. Mol Cancer Ther 15, 1427–1435.

18. Hafner, M., Niepel, M., Chung, M., and Sorger, P.K. (2016). Growth rate inhibition metrics correct for confounders in measuring sensitivity to cancer drugs. Nat Methods 13, 521–527.

19. Han, H., Jain, A.D., Truica, M.I., Izquierdo-Ferrer, J., Anker, J.F., Lysy, B., Sagar, V., Luan, Y., Chalmers, Z.R., Unno, K., et al. (2019). Small-Molecule MYC Inhibitors Suppress Tumor Growth and Enhance Immunotherapy. Cancer Cell 36, 483–497 e415.

20. Hart, T., Chandrashekhar, M., Aregger, M., Steinhart, Z., Brown, K.R., MacLeod, G., Mis, M., Zimmermann, M., Fradet-Turcotte, A., Sun, S., et al. (2015). High-Resolution CRISPR Screens Reveal Fitness Genes and Genotype-Specific Cancer Liabilities. Cell 163, 1515–1526.

21. Hayes, T.K., Neel, N.F., Hu, C., Gautam, P., Chenard, M., Long, B., Aziz, M., Kassner, M., Bryant, K.L., Pierobon, M., et al. (2016). Long-Term ERK Inhibition in KRAS-Mutant Pancreatic Cancer Is Associated with MYC Degradation and Senescence-like Growth Suppression. Cancer Cell 29, 75–89.

22. Hirabayashi, Y., Nomura, K.H., and Nomura, K. (2013). The acetyl-CoA transporter family SLC33. Mol Aspects Med 34, 586–589.

23. Huber, K.V., Salah, E., Radic, B., Gridling, M., Elkins, J.M., Stukalov, A., Jemth, A.S., Gokturk, C., Sanjiv, K., Stromberg, K., et al. (2014). Stereospecific targeting of MTH1 by (S)-crizotinib as an anticancer strategy. Nature 508, 222–227.

24. Jost, M., Santos, D.A., Saunders, R.A., Horlbeck, M.A., Hawkins, J.S., Scaria, S.M., Norman, T.M., Hussmann, J.A., Liem, C.R., Gross, C.A., et al. (2020). Titrating gene expression using libraries of systematically attenuated CRISPR guide RNAs. Nat Biotechnol 38, 355–364.

25. Kessler, J.D., Kahle, K.T., Sun, T., Meerbrey, K.L., Schlabach, M.R., Schmitt, E.M., Skinner, S.O., Xu, Q., Li, M.Z., Hartman, Z.C., et al. (2012). A SUMOylation-dependent transcriptional subprogram is required for Myc-driven tumorigenesis. Science 335, 348–353.

26. Koch, H.B., Zhang, R., Verdoodt, B., Bailey, A., Zhang, C.D., Yates, J.R., 3rd, Menssen, A., and Hermeking, H. (2007). Large-scale identification of c-MYC-associated proteins using a combined TAP/MudPIT approach. Cell Cycle 6, 205-217.

27. Lee, H.C., Wang, H., Baladandayuthapani, V., Lin, H., He, J., Jones, R.J., Kuiatse, I., Gu, D., Wang, Z., Ma, W., et al. (2017). RNA Polymerase I Inhibition with CX-5461 as a Novel Therapeutic Strategy to Target MYC in Multiple Myeloma. Br J Haematol 177, 80–94.

28. Lin, C.Y., Loven, J., Rahl, P.B., Paranal, R.M., Burge, C.B., Bradner, J.E., Lee, T.I., and Young, R.A. (2012). Transcriptional amplification in tumor cells with elevated c-Myc. Cell 151, 56–67.

29. Lin, K.H., Rutter, J.C., Xie, A., Pardieu, B., Winn, E.T., Bello, R.D., Forget, A., Itzykson, R., Ahn, Y.R., Dai, Z., et al. (2020). Using antagonistic pleiotropy to design a chemotherapy-induced evolutionary trap to target drug resistance in cancer. Nat Genet 52, 408–417.

30. Liu, L., Ulbrich, J., Muller, J., Wustefeld, T., Aeberhard, L., Kress, T.R., Muthalagu, N., Rycak, L., Rudalska, R., Moll, R., et al. (2012). Deregulated MYC expression induces dependence upon AMPK-related kinase 5. Nature 483, 608–612.

31. Luo, J., Emanuele, M.J., Li, D., Creighton, C.J., Schlabach, M.R., Westbrook, T.F., Wong, K.K., and Elledge, S.J. (2009a). A genome-wide RNAi screen identifies multiple synthetic lethal interactions with the Ras oncogene. Cell 137, 835–848.

32. Luo, J., Solimini, N.L., and Elledge, S.J. (2009b). Principles of cancer therapy: oncogene and non-oncogene addiction. Cell 136, 823–837.

33. Martin, T.D., Cook, D.R., Choi, M.Y., Li, M.Z., Haigis, K.M., and Elledge, S.J. (2017). A Role for Mitochondrial Translation in Promotion of Viability in K-Ras Mutant Cells. Cell Rep 20, 427–438.

34. Meerbrey, K.L., Hu, G., Kessler, J.D., Roarty, K., Li, M.Z., Fang, J.E., Herschkowitz, J.I., Burrows, A.E., Ciccia, A., Sun, T., et al. (2011). The pINDUCER lentiviral toolkit for inducible RNA interference in vitro and in vivo. Proc Natl Acad Sci U S A 108, 3665–3670.

35. Metallo, C.M., Gameiro, P.A., Bell, E.L., Mattaini, K.R., Yang, J., Hiller, K., Jewell, C.M., Johnson, Z.R., Irvine, D.J., Guarente, L., et al. (2011). Reductive glutamine metabolism by IDH1 mediates lipogenesis under hypoxia. Nature 481, 380–384.

36. Mycielska, M.E., Dettmer, K., Rummele, P., Schmidt, K., Prehn, C., Milenkovic, V.M., Jagla, W., Madej, G.M., Lantow, M., Schladt, M., et al. (2018). Extracellular Citrate Affects Critical Elements of Cancer Cell Metabolism and Supports Cancer Development In Vivo. Cancer Res 78, 2513–2523.

37. Panda, S., Banerjee, N., and Chatterjee, S. (2020). Solute carrier proteins and c-Myc: a strong connection in cancer progression. Drug Discov Today 25, 891–900.

38. Pavlova, N.N., and Thompson, C.B. (2016). The Emerging Hallmarks of Cancer Metabolism. Cell Metab 23, 27–47.

39. Pelossof, R., Fairchild, L., Huang, C.H., Widmer, C., Sreedharan, V.T., Sinha, N., Lai, D.Y., Guan, Y., Premsrirut, P.K., Tschaharganeh, D.F., et al. (2017). Prediction of potent shRNAs with a sequential classification algorithm. Nat Biotechnol 35, 350–353.

40. Roig, A.I., Eskiocak, U., Hight, S.K., Kim, S.B., Delgado, O., Souza, R.F., Spechler, S.J., Wright, W.E., and Shay, J.W. (2010). Immortalized epithelial cells derived from human colon biopsies express stem cell markers and differentiate in vitro. Gastroenterology 138, 1012–1021 e1011-1015.

41. Romero, R., Sánchez-Rivera, F.J., Westcott, P.M.K., Mercer, K.L., Bhutkar, A., Muir, A., Robles, T.J.G., Rodríguez, S.L., Liao, L.Z., Ng, S.R., et al. (2020). Keap1 mutation renders lung adenocarcinomas dependent on Slc33a1. Nature Cancer 1, 589–602.

42. Sack, L.M., Davoli, T., Li, M.Z., Li, Y., Xu, Q., Naxerova, K., Wooten, E.C., Bernardi, R.J., Martin, T.D., Chen, T., et al. (2018). Profound Tissue Specificity in Proliferation Control Underlies Cancer Drivers and Aneuploidy Patterns. Cell 173, 499–514 e423.

43. Schaub, F.X., Dhankani, V., Berger, A.C., Trivedi, M., Richardson, A.B., Shaw, R., Zhao, W., Zhang, X., Ventura, A., Liu, Y., et al. (2018). Pan-cancer Alterations of the MYC Oncogene and Its Proximal Network across the Cancer Genome Atlas. Cell Syst 6, 282–300 e282.

44. Sears, R., Nuckolls, F., Haura, E., Taya, Y., Tamai, K., and Nevins, J.R. (2000). Multiple Ras-dependent phosphorylation pathways regulate Myc protein stability. Genes Dev 14, 2501–2514.

45. Soucek, L., Helmer-Citterich, M., Sacco, A., Jucker, R., Cesareni, G., and Nasi, S. (1998). Design and properties of a Myc derivative that efficiently homodimerizes. Oncogene 17, 2463–2472.

46. Stewart, T.A., Pattengale, P.K., and Leder, P. (1984). Spontaneous mammary adenocarcinomas in transgenic mice that carry and express MTV/myc fusion genes. Cell 38, 627–637.

47. Stine, Z.E., Walton, Z.E., Altman, B.J., Hsieh, A.L., and Dang, C.V. (2015). MYC, Metabolism, and Cancer. Cancer Discov 5, 1024–1039.

48. Topham, C., Tighe, A., Ly, P., Bennett, A., Sloss, O., Nelson, L., Ridgway, R.A., Huels, D., Littler, S., Schandl, C., et al. (2015). MYC Is a Major Determinant of Mitotic Cell Fate. Cancer Cell 28, 129–140.

49. Toyoshima, M., Howie, H.L., Imakura, M., Walsh, R.M., Annis, J.E., Chang, A.N., Frazier, J., Chau, B.N., Loboda, A., Linsley, P.S., et al. (2012). Functional genomics identifies therapeutic targets for MYC-driven cancer. Proc Natl Acad Sci U S A 109, 9545–9550.

50. Vaseva, A.V., Blake, D.R., Gilbert, T.S.K., Ng, S., Hostetter, G., Azam, S.H., Ozkan-Dagliyan, I., Gautam, P., Bryant, K.L., Pearce, K.H., et al. (2018). KRAS Suppression-Induced Degradation of MYC Is Antagonized by a MEK5-ERK5 Compensatory Mechanism. Cancer Cell 34, 807–822 e807.

51. Wang, T., Birsoy, K., Hughes, N.W., Krupczak, K.M., Post, Y., Wei, J.J., Lander, E.S., and Sabatini, D.M. (2015). Identification and characterization of essential genes in the human genome. Science 350, 1096–1101.

52. Wanzel, M., Herold, S., and Eilers, M. (2003). Transcriptional repression by Myc. Trends Cell Biol 13, 146–150.

53. Watson, D.K., Psallidopoulos, M.C., Samuel, K.P., Dalla-Favera, R., and Papas, T.S. (1983). Nucleotide sequence analysis of human c-myc locus, chicken homologue, and myelocytomatosis virus MC29 transforming gene reveals a highly conserved gene product. Proc Natl Acad Sci U S A 80, 3642–3645.

54. Welcker, M., Orian, A., Jin, J., Grim, J.E., Harper, J.W., Eisenman, R.N., and Clurman, B.E. (2004). The Fbw7 tumor suppressor regulates glycogen synthase kinase 3 phosphorylation-dependent c-Myc protein degradation. Proc Natl Acad Sci U S A 101, 9085–9090.

55. Xiang, Z., Luo, H., Payton, J.E., Cain, J., Ley, T.J., Opferman, J.T., and Tomasson, M.H. (2010). Mcl1 haploinsufficiency protects mice from Myc-induced acute myeloid leukemia. J Clin Invest 120, 2109–2118.

56. Yada, M., Hatakeyama, S., Kamura, T., Nishiyama, M., Tsunematsu, R., Imaki, H., Ishida, N., Okumura, F., Nakayama, K., and Nakayama, K.I. (2004). Phosphorylation-dependent degradation of c-Myc is mediated by the F-box protein Fbw7. EMBO J 23, 2116–2125.

57. Yue, M., Jiang, J., Gao, P., Liu, H., and Qing, G. (2017). Oncogenic MYC Activates a Feedforward Regulatory Loop Promoting Essential Amino Acid Metabolism and Tumorigenesis. Cell Rep 21, 3819–3832.

58. Zirin, J., Ni, X., Sack, L.M., Yang-Zhou, D., Hu, Y., Brathwaite, R., Bulyk, M.L., Elledge, S.J., and Perrimon, N. (2019). Interspecies analysis of MYC targets identifies tRNA synthetases as mediators of growth and survival in MYC-overexpressing cells. Proc Natl Acad Sci U S A 116, 14614–14619.

